# Population assessment and habitat associations of the Visayan Hornbill *Penelopides panini* in Northwest Panay, Philippines

**DOI:** 10.1101/2021.08.31.458333

**Authors:** Holly Isabelle Mynott, David Charles Lee, Rhea Aranas Santillan, Christian Jürgen Schwarz, Benjamin Tacud, Arcel Dryden Fernandez, Daphne Kerhoas

## Abstract

**Background:** Seven out of 10 hornbill species in the Philippines are threatened with extinction. Among these is the Endangered Visayan hornbill *Penelopides panini*, found on Panay and Negros islands. Threatened by habitat loss and hunting, its population size is thought to have declined from 1,800 individuals 20 years ago to less than 1,000. However, a recent study on Negros estimated 3,564 individuals across three core forest blocks. This study aims to quantify the Visayan hornbill population size in and around the Northwest Panay Peninsula Natural Park (NWPPNP) on Panay, the largest contiguous low-elevation forest landscape remaining across its range, and its broad habitat associations across a gradient of environmental degradation.

**Methods:** Hornbills were surveyed using 10-minute Distance sampling point counts (n = 362) along transects (average length 1.1 km). Habitat variables were recorded, while habitat was classified into: primary forest, secondary forest, plantation or open habitat. Using Distance software, population densities were estimated for, and post-stratified by habitat, with the overall population estimate taken as a mean of habitat density estimates weighted by habitat area. Using logistic binary regression, hornbill occurrence was modelled against reduced habitat factors extracted from factor analysis of the habitat data.

**Results:** Surveys covered 204.4 km^2^ of the 374.8 km^2^ Northwest Panay Peninsula. Hornbills were not recorded in plantation or open habitats. The estimated density of hornbills was significantly higher in primary forest (17.7 individuals km^−2^ ± 29.7% CV) than in secondary forest (5.0 individuals km^−2^ ± 36.7 %CV; *z* = 9.538, *P* < 0.001). The overall population estimate is 2,231 individuals ± 24.4 %CV for the NWPPNP and environs, and 2,949 individuals ± 23.1 %CV for the entire Northwest Panay Peninsula. One habitat factor, described by increasing numbers of large trees, elevation and distance from the Park’s boundary, had a significant positive effect in explaining hornbill occurrence, with hornbills significantly more likely to occur in primary forest than the other habitat types.

**Conclusions:** Our study demonstrates the habitat preference of the Visayan hornbill, highlights the importance of the NWPPNP for the species’ conservation, and provides strong evidence for re-assessing the global population size.

## Background

The Philippines is one of the 18 mega-biodiverse countries of the world, a collective which harbours two thirds of the earth’s biodiversity, while the Philippines itself ranks fourth in levels of global bird endemism (Convention on Biological Diversity 2021). However, Southeast Asia as a whole is experiencing a wildlife crisis (Harrison et al. 2016), primarily due to some of the highest deforestation rates in the world (Hughes, 2017), and severe hunting pressures (Gray et al. 2018).

Hornbills, as frugivorous birds, play an important role in seed dispersal in tropical forests (Kinnaird and O’Brien 2007). Targeted for domestic consumption and/or the international trade in their casques (Sreekar et al. 2015), their tendency to congregate at fruiting trees and travel long distances makes them particularly vulnerable to hunting pressure (Harrison et al. 2016). Furthermore, as cavity-nesting species, they are also vulnerable to deforestation, as they often rely on old, larger trees in undisturbed forest to breed (Kinnaird and O’Brien 2007). As a result, seven out of the 10 hornbill species in the Philippines are considered globally threatened with extinction, among which four are Endangered or Critically Endangered, while the populations of all 10 species are thought to be decreasing (IUCN 2021).

Endemic to the Western Visayas of the Philippines (Collar et al. 1999), the globally Endangered Visayan (or Tarictic) hornbill (*Penelopides panini*) is recorded on the islands of Negros and Panay. Small populations may remain on Masbate and Pan de Azucar (BirdLife International 2020), while it is now considered locally extinct on Ticao (Curio, 1994; del Hoyo et al. 2001), Guimaras and Sicogon (BirdLife International 2020). The remaining populations exhibit loss of genetic diversity and are probably genetically isolated due to at least 100 km distance between currently available habitats (Sammler et al. 2012). The Visayan hornbill inhabits dipterocarp forest up to 1,100 m, occasionally to 1,500 m, and with a preference for undisturbed habitat, although it does utilise secondary forests (BirdLife International 2020). Studies on diet and food provisioning were carried out by Klop et al. (1999) and Luft et al. (2002), while the breeding biology of Panay populations was studied by Klop et al. (2000) and Curio (2006).

As with other hornbills in the region, the Visayan hornbill is threatened by deforestation and hunting, resulting in an increasingly fragmented, small population (BirdLife International 2020). While natural forest cover (>30% canopy cover, vegetation >5 m height) was estimated at ~37% and ~27% on Panay and Negros, respectively, in 2000, only 5.4% and 2.1% of this was classified as primary forest (Global Forest Watch 2021). Very small forest fragments remain on the species’ other range islands (BirdLife International 2020). Previously estimated at 1,800+ individuals, and 1,200 mature individuals (BirdLife International 2001), declines in the last 20 years suggest the global population may now comprise <1,000 individuals, with no subpopulation containing more than 250 individuals (BirdLife International 2020).

Despite global pressures on hornbills, surveys of the Visayan hornbill have suggested larger population sizes than were previously estimated. A study around Mt. Balabac on Panay found an average density of three nests km^−2^, and extrapolated the total breeding population on Panay to be in the range of 750 to 1,500 pairs, under the assumption of 225-450 km^2^ of suitable habitat remaining (Klop et al. 2000). A 6-year study by the Philippine Biodiversity Conservation Foundation estimated a population of 3,564 individuals across three forest blocks in Negros island (Chavez 2020). This island estimate was calculated using a distance sampling point count method, and based on an overall lowland forest density of 14 individuals km^−2^. Both these numbers are considerably greater than the last global population estimate for the species (BirdLife International 2020), which is encouraging for the species’ conservation.

This study aims to complement the population studies on Negros and the Central Panay Mountain Range by quantifying the Visayan hornbill’s population size within the Northwest Panay Peninsula Natural Park (NWPPNP) and surrounding peninsula on Panay. At 120 km^2^, NWPPNP is the largest remaining contiguous low-elevation forest landscape remaining across the hornbill’s range (BirdLife International 2021a), and where it is reported to be “common” (Curio and Schwarz 2017). This study quantifies the Visayan hornbill’s population size in this key landscape for the conservation of this threatened species, and generates habitat-specific density estimates, contributing robust data to inform and revise the current global population estimate. It also quantifies the hornbill’s habitat associations across a gradient of environmental degradation, from primary forest to open habitat.

## Methods

### Study site

This survey took place in the Northwest Panay Peninsula Natural Park (NWPPNP; Figure 1; 122.0003, 11.8130), a protected area () since 2002 (PhilinCon 2021). Under the National Integrated Protected Areas System (NIPAS) Act, Philippines, Natural Parks (comparable to IUCN Category II Protected Areas, National Parks) are landscapes not altered significantly by anthropic activity, and managed to maintain their natural, national or international significance (La Viña et al. 2010). However, while most resource extraction is prohibited, development which is considered ‘sustainable,’ such as renewable energy generation projects, may now be permitted inside buffer zones within park borders (Congress of the Philippines 2018). The NWPPNP is also an Important Bird and Biodiversity Area (IBA; BirdLife International 2021a), and Key Biodiversity Area (KBA; Key Biodiversity Area Partnership, 2020), for which the Visayan hornbill, as a geographically restricted species, is one of five bird species triggering the site’s KBA classification (Key Biodiversity Area Partnership, 2020). The Park (27 to 875 m elevation) covers 120 km^2^ of tall dipterocarp, limestone karst, lower montane, and bamboo forests, including 25-50 km^2^ of old growth tropical forest (BirdLife International 2021a). It is likely to be the largest remaining area of contiguous lowland forest in the Negros and Panay Endemic Bird Area (Key Biodiversity Area Partnership, 2020).

**Figure 1:**
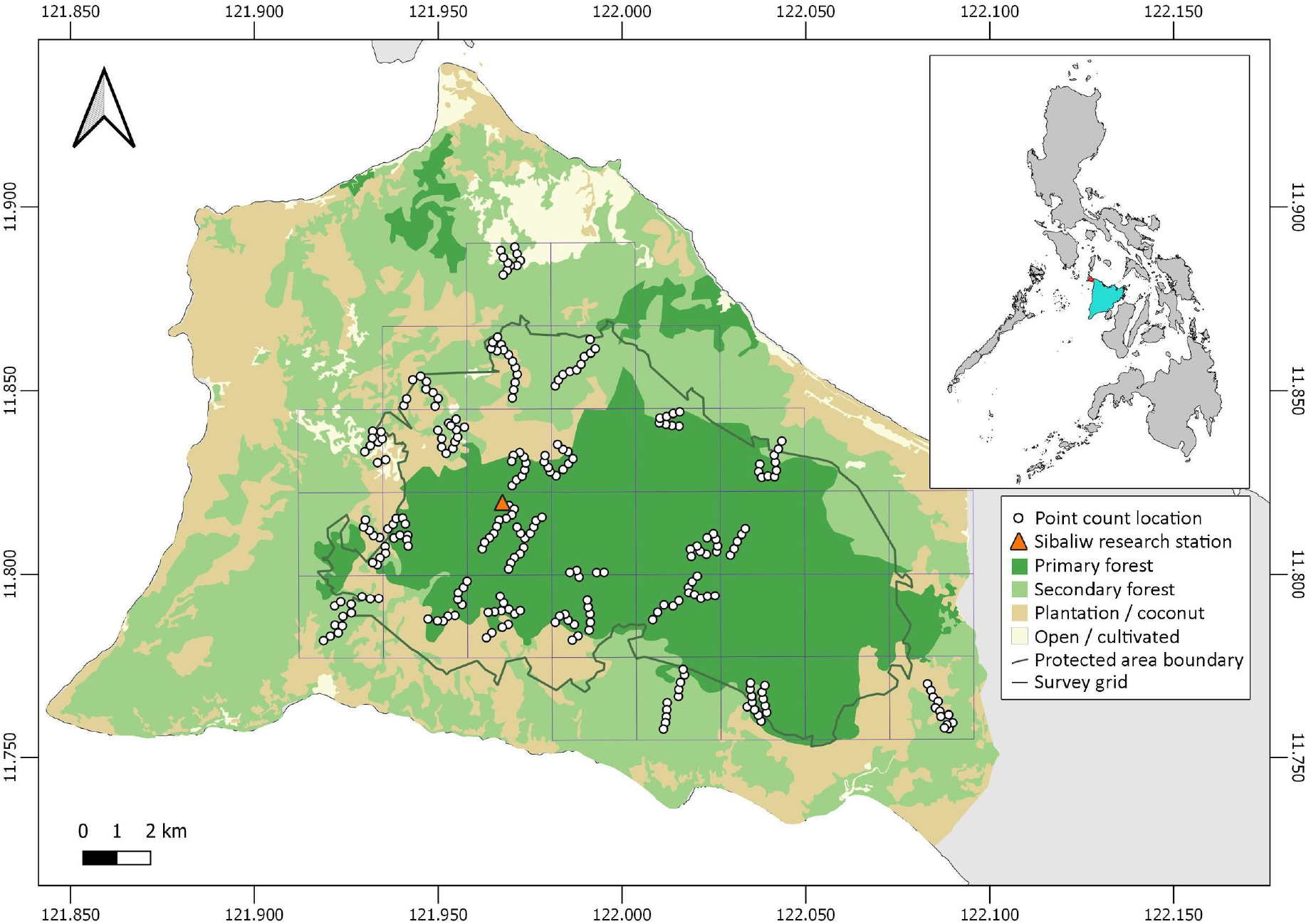
Map of the survey site and its location within the Northwest Panay Peninsula, Panay, Philippines.

While the human population within the protected area is relatively low (BirdLife International 2021a), there are many settlements located at the forest edge (e.g. La Serna, Paua, Tagosip, Codiong), with some farming legally ongoing inside the park on plots that existed prior to its establishment (Santillan, R.A. *pers. obs*., 2019), in addition to illegal plots established after the creation of the Park. The Natural Park is managed by the municipal government, the local Department of Environment and Natural Resources (DENR) under NIPAS, and non-governmental organisations (Mogul and Aquino-Ong 2016).

Threats to biodiversity in the NWPPNP include illegal logging and land conversion for slash-and-burn agriculture (kaingin), with some natural forest converted to plantation, and hunting, which is thought to have significantly impacted several bird and mammal species (BirdLife International 2021a). Mining applications encompass the remaining forest cover of the protected area (Key Biodiversity Area Partnership 2020; PhilinCon 2021).

### Survey method

The study site was overlaid with a grid of 2.5 x 2.5 km cells (n = 33), all of which included at least some of the NWPPNP (Figure 1). Any cell that included the shore was removed, as the human population is concentrated in these areas and the habitat is mainly heavily disturbed. Using a random selection function, 24 cells were identified to survey (72.7% of the study area). However, once on site, three of the selected cells were found to be inaccessible due to very steep terrain. In these instances, the nearest accessible grid cell was surveyed instead. All 24 cells were surveyed once, ,with surveys repeated in seven cells (29.2%), before access restrictions in March 2020 in response to the coronavirus pandemic halted further repeats. Within each grid cell, 1 to 4 transects of 0.3 to 2.5 km 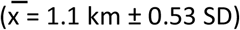 were positioned randomly, or along existing narrow trails if the terrain was particularly difficult. Since any trails used were <1 m width, this positioning of effort is considered not to have biased results (Cornils et al. 2015). Overall transect effort, not including repeats, was 1 to 3.5 km cell^−1^ 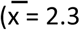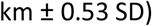.

Distance sampling point counts (Lloyd et al. 2000; Lee and Marsden, 2008) were used to survey hornbills along transects. This approach makes a number of critical assumptions, which the survey was designed to meet: (1) transect lines are randomly placed with respect to species distribution; (2) birds directly on the transect are always detected; (3) birds are detected at their original location before movement; and (4) distances are accurately measured (Buckland et al. 2001). Point counts were situated ≥200 m apart along transects, minimising the likelihood of recording the same birds from multiple points (Bibby et al. 1998; Marsden 1999). In total, 362 point counts were surveyed, including 84 repeated counts.

A count period of 10 minutes was used as a compromise between maximizing detection and minimising multiple-counting, and as used previously for hornbill surveys (Marsden 1999). Radial distances to birds were estimated using laser rangefinders (Nikon Aculon AL11 and Volvik V1 models) whenever possible. If a clear line of sight was unobtainable, observers then estimated radial distance. Pre-survey training in distance estimationensured there was no between-observer bias in consistently over- or under-estimating distances (Bibby et al. 1998). Point counts were conducted throughout the day, from 08:00 to 18:00; long morning and afternoon hours have been used in previous hornbill surveys (Marsden and Pilgrim, 2003; Naniwadekar and Datta, 2013), and previous work in the study site has recorded hornbill activity throughout the day (Klop et al. 2000; Schwarz, C.J. *pers. obs*., 2019). Point counts were not carried out in heavy rain or high winds (Marsden and Pilgrim, 2003).

### Habitat surveys

The habitat at each point count location was classified into one of four categories: primary forest (old-growth forest), secondary forest, plantation or open habitat. Forests were classified as primary or secondary based on structural appearance and floristic composition, known history and local knowledge, and supported by satellite imagery (Stouffer et al. 2006). Plantation forest was mostly areas of palm trees or young rainforest interspersed with palms, with the occasional young mahogany plantation. Open habitats included areas of scrub or low-lying ferns with few trees.

Habitat categorisation across the Park was supported by plot-based surveys every 500 m along transects (n = 160 plots). Within plots of 20 m diameter (0.03 ha), numbers of tree stems with a diameter at breast height (DBH) of 10 ≤25 cm, >25 ≤99 cm, and ≥100 cm were recorded, the latter based on the hornbill’s ecological requirement of large trees for nesting (Klop et al. 2000). The branching architecture of any tree with ≥100 cm DBH was described, following Bibby et al. (1998)an indication of forest (disturbance) history; closed canopy, open canopy, or regenerating forest habitats. Canopy cover was calculated from four readings (north, east, south and west facing) from the centre of each plot using a spherical crown densiometer (CSP Forestry concave model). The elevation of each plot and its distance from the Park’s boundary was recorded, the latter as a proxy for accessibility and potential anthropic disturbance.

### Data analysis

To categorise habitat across the remaining peninsula outside the survey area, a dataset of land cover was used (PhilGIS 2020), which categorised land into 13 categories, from closed canopy primary forest to built-up land. These areas were re-categorised into the same four categories as measured in our survey, based on mapping our measured habitat variables against the PhilGIS dataset, using QGIS version 3 (QGIS Development Team 2020) to compare datasets.

Hornbill survey data were analysed using conventional distance sampling in Distance software version 7.3 (Thomas et al. 2010), which models the reduction in detection probability of the focal animal as (radial) distance from the observer increases (Buckland et al. 2001). Where group size was uncertain, a habitat-specific mean group size taken from visual detections was used (Lee and Marsden 2008). After preliminary data exploration, data were grouped into 20 m distance bins and truncated at 60 m. A series of detection functions (uniform, half-normal, hazard rate) and expansion terms (cosine, simple and hermite polynomials) were applied to the truncated data, sequentially. Population densities were estimated for, and post-stratified by habitat, with the overall population estimate taken as a mean of habitat density estimates weighted by habitat area. Final model selection was based on AIC values and goodness of fit statistics (Buckland et al. 1993). Two-sample *z*-tests were used to analyse differences between habitat-specific density estimates computed by Distance (Thomas et al. 2010), and using the formula:

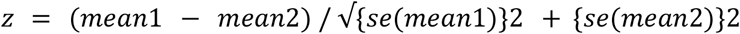

Factor Analysis with principal components extraction (PCA) and varimax rotation was carried out to reduce the nine original habitat variables into a smaller number of extracted habitat factors and correct for any collinearity between variables. Only factors with eigenvalues of >1.0 were considered in the final analysis. Binary logistic regression was then used to model the occurrence of hornbill at the plot level against all PCA factors and their interaction covariates. Following that, a final regression model was refitted, including only those covariates with significant model coefficients. A Hosmer-Lemeshow goodness-of-fit test (Hosmer and Lemeshow 2000) was used to assess the fit of the final model, and a one-way analysis of variance was used to test for differences in factor scores across habitat types.

## Results

In total, surveys of the NWPPNP and environs covered 204.4 km^2^ (54.5%) of the 374.8 km^2^ Northwest Panay Peninsula. Within the peninsula, delimited on the east by the road from Kalibo and Caticlan to Pandan (easternmost point: 121.1032, 11.7503), about 106 km^2^ was classified as primary forest, 156 km^2^ as secondary forest, 101 km^2^ as plantation and 16 km^2^ as open area. The Visayan hornbill was recorded 70 times, in 16 of the 24 grid cells; a landscape naive occurrence of 67.7%. Hornbills were not recorded in plantation or open habitats.

Based on the data collected through remote spatial resources and habitat plots, primary forest covers about 70% of the protected area (91 km^2^), secondary forest 16% (21 km^2^) and other areas 14% (19 km^2^), most of which is coconut plantation. Of the 160 habitat plots, 77 were in the primary forest, 49 in secondary forest, 22 in plantations and 12 in open habitat.

### Density and population estimates

There were no sightings of Visayan hornbills in plantation areas or open habitats. Therefore, distance data were only modelled for primary and secondary forest habitats. In both cases, a uniform detection function best fitted the data, and with a simple polynomial or cosine adjustment term for primary and secondary forest, respectively (Figure 2). The estimated density of hornbills was significantly higher in primary forest (17.7 individuals km^−2^ ± 29.7 %CV) than in secondary forest (5.0 individuals km^−2^ ± 36.7 %CV; *z* = 9.538, *P* < 0.001). Using these densities generated habitat-specific population estimates of 1,729 individuals ± 29.7 %CV (972 - 3,077 95% CIs) in primary forest and 501 individuals ± 36.7 %CV (248 - 1,014 95% CIs) in secondary forest. The overall population estimate, weighted by habitat area and pooled across all habitat strata, for the NWPPNP and its immediate surroundings is 2,231 ± 24.4 %CV (1,385 - 3,592 95% CIs). For the entire Northwest Panay Peninsula (374.8 km^2^), the hornbill population is estimated at 2,949 individuals ± 23.1 %CV; 1,852 individuals ± 29.7 %CV in primary forest and 1,097 ± 36.7 %CV in secondary forest (Table 1).

**Table 1:**
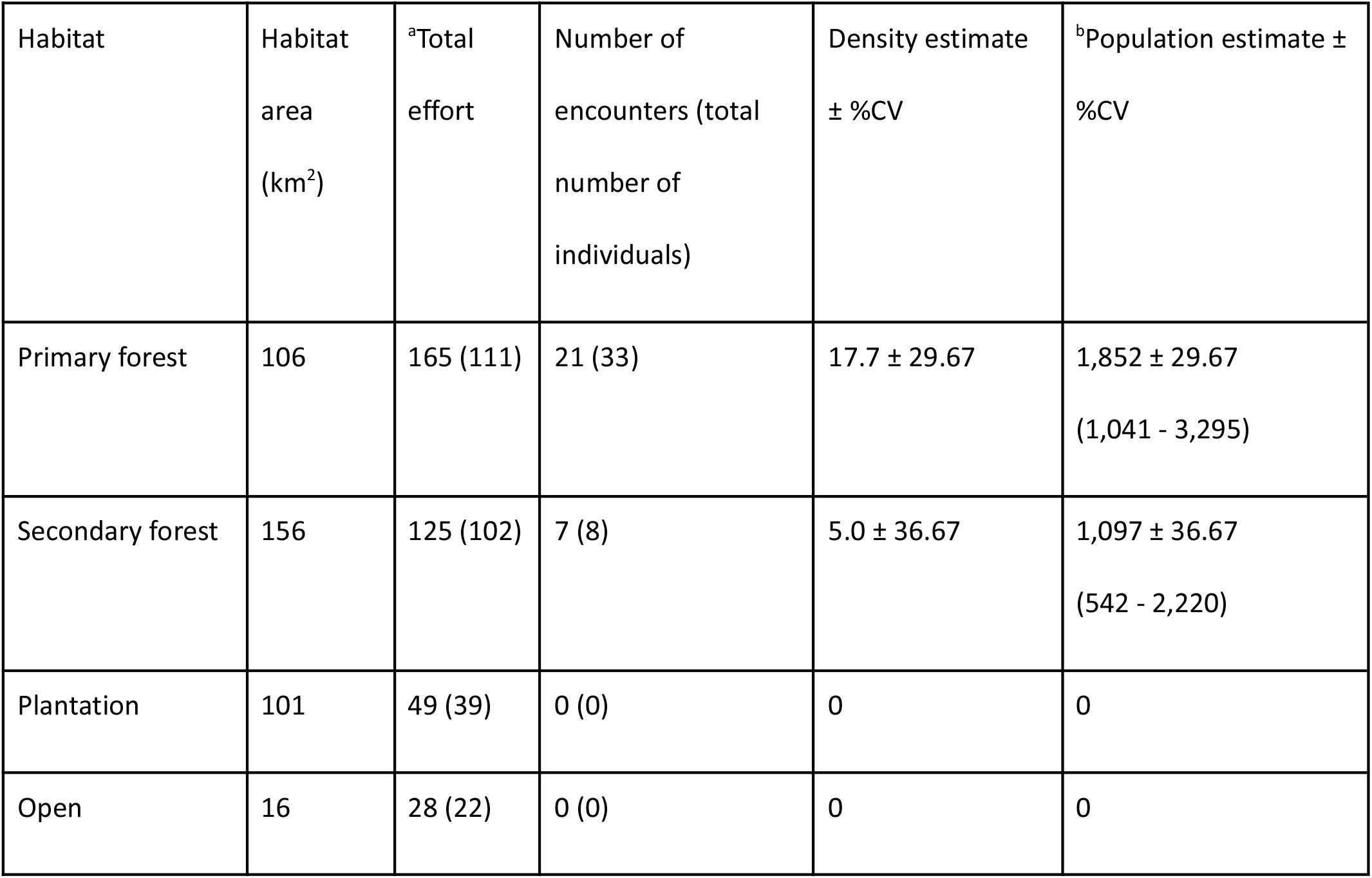
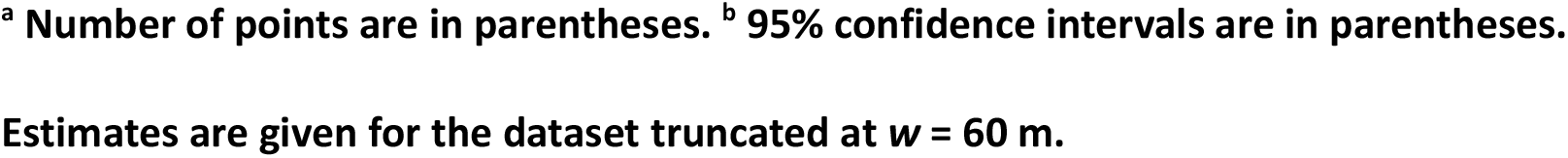
Habitat-specific density and population estimates for the Northwest Panay Peninsula.

| Habitat | Habitat area (km<sup>2</sup>) | <sup>a</sup>Total effort | Number of encounters (total number of individuals) | Density estimate ± %CV | <sup>b</sup>Population estimate ± %CV |
| --- | --- | --- | --- | --- | --- |
| Primary forest | 106 | 165 (111) | 21 (33) | 17.7 ± 29.67 | 1,852 ± 29.67<br>(1,041 - 3,295) |
| Secondary forest | 156 | 125 (102) | 7 (8) | 5.0 ± 36.67 | 1,097 ± 36.67<br>(542 - 2,220) |
| Plantation | 101 | 49 (39) | 0 (0) | 0 | 0 |
| Open | 16 | 28 (22) | 0 (0) | 0 | 0 |

<sup>a</sup> Number of points are in parentheses. <sup>b</sup> 95% confidence intervals are in parentheses.
Estimates are given for the dataset truncated at $w = 60$ m.

**Figure 2.**
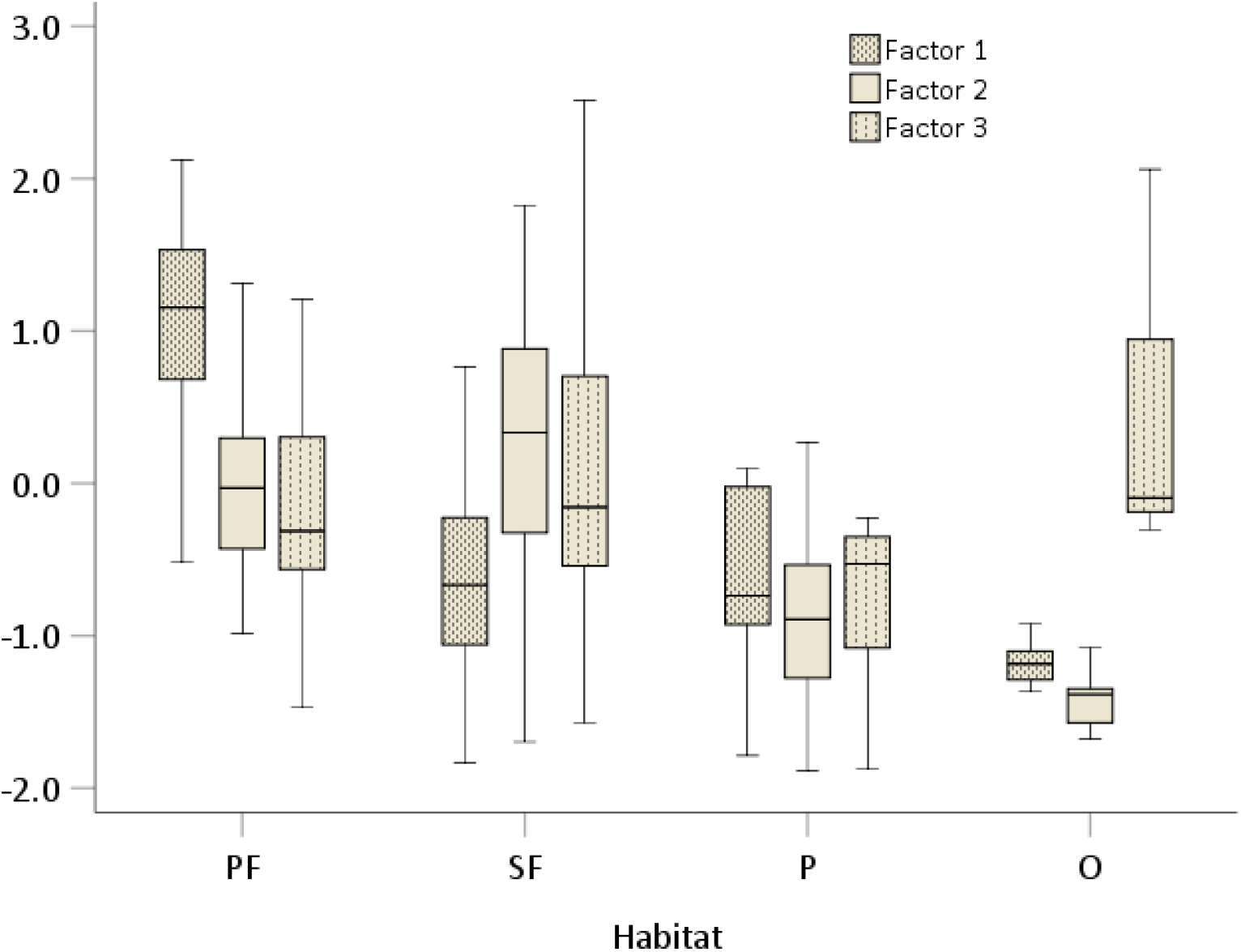
Boxplots of median and interquartile ranges of PCA Factor scores for each habitat type; Primary forest (PF), Secondary forest (SF), Plantation (P), and Open (O).

### Habitat associations

Three PCA factors explained 62.3% of the variance in the nine original habitat variables (Table 2; Figure 2). Factor 1 (27.5% of the explained variance in the original habitat variables) was associated with increasing numbers of large tree stems indicative of a closed canopy forest history, increasing elevation and distance from the Park’s boundary, and decreasing prevalence of trees indicative of closing canopy or regenerating forest histories. Factor 2 (21.9%) explained a habitat gradient of increasing numbers of tree stems of all size classes, but particularly trees <100 cm DBH, and increasing elevation, canopy cover, and trees indicating a regenerating forest history. Factor 3 (12.9%) described a gradient of decreasing habitat recovery, with tree structures characteristic of past open canopy forest alongside a decreasing prevalence of trees indicative of a closed canopy or regenerating forest history. Factor 1 scores were significantly higher for both primary and secondary forest than plantation and open habitat, and higher for primary forest than secondary forest (*F*_3,86_ = 32.000, *P* <0.001, with Bonferroni post hoc test). Factor 2 scores were significantly higher for both primary and secondary forest than plantation and open habitat (*F*_3,86_ = 17.636, *P* <0.001, with Bonferroni post hoc test).

**Table 2.** Rotated component matrix of a PCA on the nine original habitat variables.

| Variable | Factor 1<br>(2.476; 27.5%) | Factor 2<br>(1.967; 21.9%) | Factor 3<br>(1.159; 12.9%) |
| --- | --- | --- | --- |
| Distance to Park boundary | +0.830 | +0.204 |  |
| Proportion of branching type A | +0.810 |  | -0.271 |
| Elevation | +0.672 | +0.304 |  |
| Trees of DBH $\geq 100$ cm | +0.628 | +0.274 | |
| Canopy cover | +0.241 | +0.640 |  |
| Trees of DBH 10-25 cm | +0.223 | +0.639 | +0.242 |
| Trees of DBH 26-99 cm |  | +0.756 |  |
| Proportion of branching types C and D | -0.380 | +0.573 | -0.484 |
| Proportion of branching type B |  |  | +0.850 |

**Only coefficients $\geq 0.2$ are displayed. Rotated eigenvalues and percentages of explained variance for each factor are in parentheses.**

The final logistic regression model included one covariate, Factor 1, which, as a predictor of hornbill presence, fitted the data adequately (Hosmer-Lemeshow test: *Ĉ*_8_ = 8.947, *P* = 0.347). The model correctly predicted hornbill presence-absence in 83.3% of cases: 94.1% correct for absence; and 50.0% correct for presence. Factor 1 had a significant positive effect on explaining the probability of hornbills utilising the habitat (*ß* = +1.233 ± 0.299, *z*^2^ = 17.052, *P* <0.001) and an odds ratio (*Exp(B)*) of 3.432 (1.911–6.162 95% CIs): for a one-unit increase in Factor 1 score, there was a 3.4 times increase in the likelihood of hornbills being present in the habitat (Figure 3). Based on the Factor 1 scores, the likelihood of hornbills occurring in primary forest (mean probability = 0.48 ± 0.038) was significantly higher than in secondary forest (0.15 ± 0.021), plantation (0.13 ± 0.023) and open habitat (0.08 ± 0.014; *H*_3_ = 43.001, *P* <0.001; pairwise comparisons with a Bonferroni correction).

**Figure 3.**
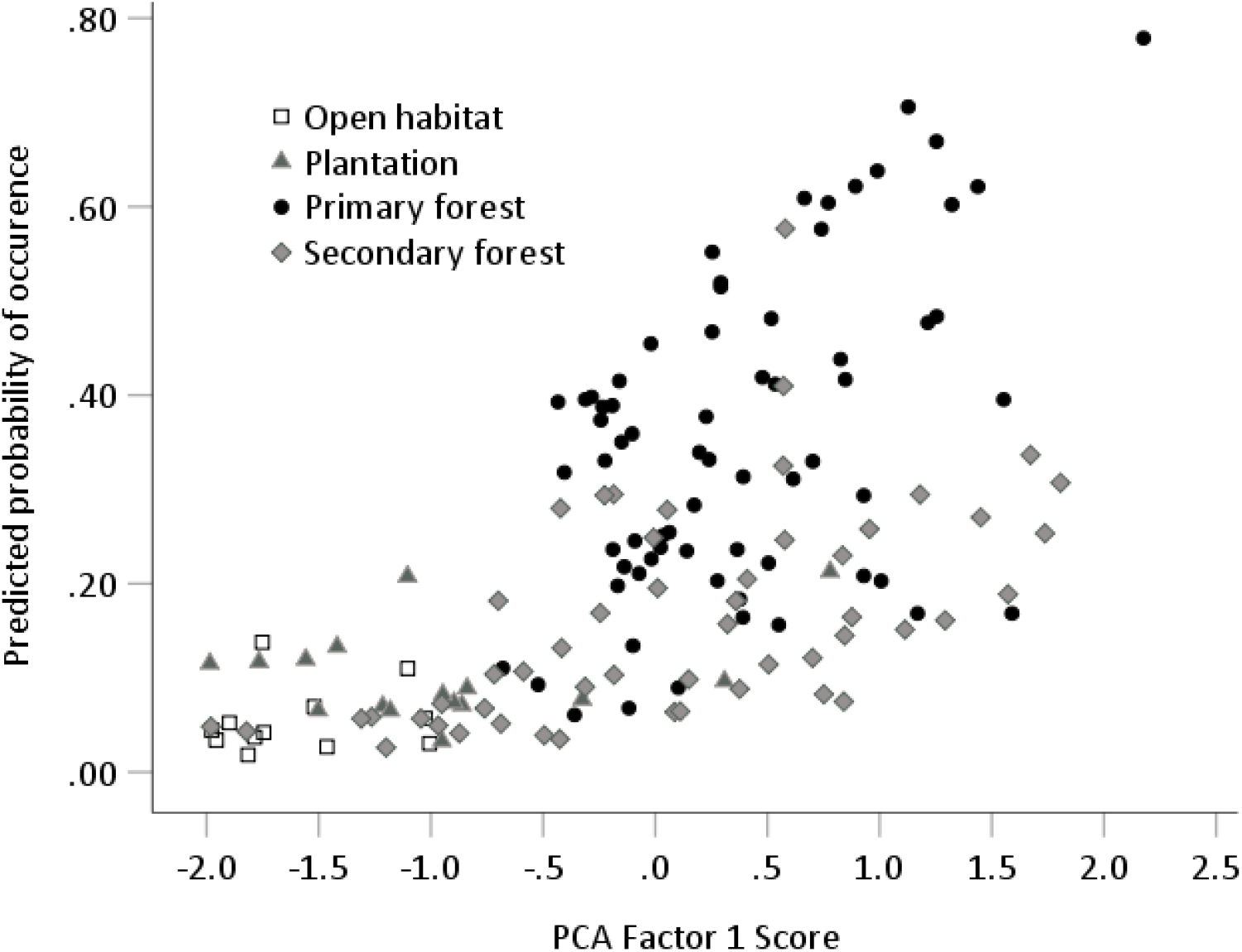
Predicted probability of occurrence of Tarictic hornbill along a habitat gradient described by PCA Factor 1.

## Discussion

The estimated population size of 2,949 within the whole Northwest Panay Peninsula is appreciably greater than the 2001 IUCN population estimate of 1,800 individuals across the species’ global range.is supported by the previous data gathered on Panay (750-1,500 breeding pairs; Klop et al. 2000), and by the Philippine Biodiversity Conservation Foundation’s recent estimate of 3,564 hornbills across three forest blocks in Negros (Chavez 2020): 1,580 individuals in North Negros Natural Park (708.3 km^2^); 532 in Mount Kanla-on Natural Park (245.6 km^2^); and 1,452 in Balinsasayao Twin Lakes Natural Park (80.2 km^2^; 290 - 1,762 m elevation; Chavez 2020). The overall lowland forest density estimate of 14 individuals km^−2^ (cited in BirdLife International 2020) from the Negros study is commensurate with that estimated in primary forest of the Northwest Panay Peninsula. Our estimate for the NWPPNP exceeds these single landscape population estimates, suggesting that the area could be a particular stronghold of this species.

The population estimates for different habitat strata within this study and the logistic regression results suggest that forest cover with minimal disturbance is highly important to the Visayan hornbill, with decreasing likelihood of occurrence in more disturbed areas. In this study, the hornbill was significantly more likely to occur in primary forest than secondary, and it was not recorded in plantation or open habitat at all. The population estimate in primary forest was also greater than that for secondary forest. This is supported by the different estimated population sizes of this species within Natural Parks on Negros. While the largest population was recorded in the geographically largest area, North Negros Natural Park, encounter rates there were lower than in Mt Kanla-on. Within North Negros Natural Park, only ~20% is forested, with some old-growth (BirdLife International 2021b). In comparison, Mt Kanla-on Natural Park has a smaller area, but about half is forested (BirdLife International 2021c). It is unclear from the Negros study how many sightings were recorded within primary or secondary forest.

As found in our study, the lower population density within North Negros Natural Park was thought to be because the park has fewer tall, mature trees suitable for hornbills (Chavez 2020). This may also explain why there is such a relatively large population in the NWPPNP, where a high density of large and mature trees remains, based on our habitat survey. It is notable that logistic regression identified no significant increase in the likelihood of hornbill presence in secondary forest compared to plantation habitat in our study. While our survey detected no Visayan hornbill in plantation forest, the PCA results suggest that secondary and plantation forest are structurally similar, based on the nine original habitat variables. Therefore, it appears the model had difficulty accurately distinguishing between these two habitats and predicting hornbill presence in secondary forests. In the absence of information on specific food resources (e.g. Naniwadekar et al. 2015), this does not confirm similar ecological value of secondary and plantation forests for Visayan hornbill. The only significant increase in the likelihood of hornbills being present was seen for primary forest, further emphasising the importance of this undisturbed habitat.

Other hornbill species have shown similar responses to reduced forest quality. In reserve forests of Arunachal Pradesh, five hornbill species and the fruiting trees they use were found at reduced abundance in heavily disturbed forests (Naniwadekar et al. 2015). In urban-forest mosaics of KwaZulu-Natal, occupancy modelling showed that large trees had a positive influence on trumpeter hornbill (*Bycanistes bucinator*) presence, while human presence negatively influenced its detection probability (Chibesa and Downs, 2017). Holbech et al. (2018) identified two small-bodied hornbill species in Ghana that are not subject to hunting pressure that have seen population declines, suggesting limited resilience to forest degradation (Holbech et al. 2018). In Ghana, while the versatile West African pied hornbill (*Lophoceros semifasciatus*) persists in fragmented forests, this fragmentation was thought to be a factor driving significant population declines (Holbech et al. 2018).

Even when hornbills are able to feed successfully in disturbed forest, they require large tree cavities for nesting (Klop et al. 2000; Marsden and Pilgrim 2003; Espanola et al. 2016). Consequently, they often breed at higher densities in primary forests (Espanola et al. 2016) and, as long-lived species, can show considerable time-lags in population decline following forest disturbance (Marsden and Pilgrim 2003). This reliance on large trees in primary forests is supported by our habitat data, in which the proportion of trees with a branching structure indicative of past old-growth forest and numbers of large trees were two of the largest contributing factors to PCA Factor 1, which significantly explained hornbill presence.

Despite protection of the NWPPNP and its large primary forest area, illegal logging still occurs (PhilinCon 2020), and the park lost 2% forest cover between 2000 and 2018 (Abrahams 2020). However, forest loss has been worse in unprotected areas over the Visayan hornbill extant range, with 4.6% of unprotected forest cover across Panay and Negros lost within the same time period (Abrahams 2020). Alongside illegal threats, the park is currently threatened by a 0.25 km^2^ hydropower development within its borders at Malay, Aklan (Antique Union for Conservation 2020). Therefore, future conservation efforts need to target and protect this natural park, a key stronghold for this species, and specifically the primary forest within this park.

A further factor affecting Visayan hornbill populations is hunting (BirdLife International 2020). Several studies have suggested that hunting may have overtaken deforestation as the greatest threat for bird species across Southeast Asia (e.g. Sreekar et al. 2015; Harrison et al. 2016). Although the NWPPNP is a protected area, the Visayan hornbill is still hunted inside the park (PhilinCon, 2020). In fact, signs of hornbill hunting (e.g. plucked feathers), rare before the coronavirus pandemic, have been witnessed since its start in 2020 (Santillan, R.A. *pers. obs*., 2020). For some species, hunting for the international wildlife trade has a critical impact, but many more are pressured by domestic consumption (Sreekar et al. 2015). In 1998, Klop et al. (2000) recorded hunting of the Visayan hornbill in the Northwest Panay Peninsula for subsistence and capture for the pet trade. Also, legs, feathers and beaks of Visayan hornbills are converted into tourist souvenirs sold on nearby Boracay Island (Schwarz, C.J. *pers. obs*., 2015). Hunting pressure may explain cases in which hornbills do not utilise certain forests or fragmented habitats (Holbech et al. 2018) and plantations, despite such disturbed areas containing suitable food resources (Marsden and Pilgrim 2003). This idea is supported by conversations with local people around the NWPPNP, who suggest that poaching pressure is the reason that hornbills, which used to visit the villages in the day time, no longer do so (Schwarz, C.J. *pers. obs*., 2019). Supporting this further, increasing distance from the park boundary (and therefore distance from human settlements and sources of hunting pressure) was also one of the largest contributing variables to PCA Factor 1, which significantly explained hornbill presence. Therefore, hunting pressure may explain why hornbills were only observed in primary and secondary forest in this study. A recent study in Mindanao, Philippines, suggests that while wildlife was traditionally hunted for sustenance, the more recent drivers of hunting within protected areas are poverty and lack of long-term livelihood options (Tanaglo 2017). Further research is recommended on the drivers for Visayan hornbill hunting within the NWPPNP.

## Conclusions

This study contributes empirically to the conservation assessment of this threatened species, providing strong evidence for re-assessing the global population size, and alongside the previous population estimates from Panay and Negros (see Klop et al. 2000; Chavez 2020). It also provides strong evidence that the Northwest Panay Peninsula, especially the protection of its Natural Park, supports a large population concentrated outside of the other significant tract of forest on Panay, the Central Panay Mountain Range (Klop et al. 2000; BirdLife International 2020), where it is considered, subjectively, fairly common (Alabado et al. 2009). As well, this study presents evidence of the species’ habitat preferences, which should guide future landscape management within and beyond the protected area. Finally, these population data provide a robust baseline against which long-term monitoring can be measured, aligning with the proposed conservation action of replicating a hornbill monitoring scheme for Panay (BirdLife International 2020).

## Declarations

### Ethics approval and consent to participate

Not applicable.

### Consent for publication

Not applicable.

### Availability of data and materials

The datasets supporting the conclusions of this article are available in the Open Science Framework repository, DOI 10.17605/OSF.IO/J9AET, at https://osf.io/j9aet/.

### Authors’ contributions

HM and DL wrote the main manuscript and analysed the hornbill population data, with HM completing the first draft and DL undertaking distance modelling according to habitat strata. DL completed the PCA and logistic regression analyses. DK organised the project design with input from DL, managed the project throughout, secured grants and other funding, and contributed significantly to editing the manuscript. RS aided logistics and permissions for survey work within the park. HM, BT and AF carried out the survey alongside others (see acknowledgements). CS provided editorial input and co-ordinated with RS for survey logistics as part of PhilinCon. All authors read and approved the final manuscript.

## Acknowledgements

We would like to thank the Philippine Department Environment of Natural Resources, and especially Mr Andres Untal (PENRO) and Mme Cynthia Blancia (CENRO) and their staff. Special thanks for the data collection to the dedicated Ilke Geladi and the rest of the team.

